# Cancer Type-Dependent Correlations between *TP53* Mutations and Antitumor Immunity

**DOI:** 10.1101/692715

**Authors:** Lin Li, Mengyuan Li, Xiaosheng Wang

## Abstract

Many studies have shown that *TP53* mutations play a negative role in antitumor immunity. However, a few studies reported that *TP53* mutations could promote antitumor immunity. To explain these contradictory findings, we analyzed five cancer cohorts from The Cancer Genome Atlas (TCGA) project. We found that *TP53*-mutated cancers had significantly higher levels of antitumor immune signatures than *TP53*-wildtype cancers in breast invasive carcinoma (BRCA) and lung adenocarcinoma (LUAD). In contrast, *TP53*-mutated cancers had significantly lower antitumor immune signature levels than *TP53*-wildtype cancers in stomach adenocarcinoma (STAD), colon adenocarcinoma (COAD), and head and neck squamous cell carcinoma (HNSC). Moreover, *TP53*-mutated cancers likely had higher tumor mutation burden (TMB) and tumor aneuploidy level (TAL) than *TP53*-wildtype cancers. However, the TMB differences were more marked between *TP53*-mutated and *TP53*-wildtype cancers than the TAL differences in BRCA and LUAD, and the TAL differences were more significant in STAD and COAD. Furthermore, we showed that TMB and TAL had a positive and a negative correlation with antitumor immunity and that TMB affected antitumor immunity more greatly than TAL did in BRCA and LUAD while TAL affected antitumor immunity more strongly than TMB in STAD and HNSC. These findings indicate that the distinct correlations between *TP53* mutations and antitumor immunity in different cancer types are a consequence of the joint effect of the altered TMB and TAL caused by *TP53* mutations on tumor immunity. Our data suggest that the *TP53* mutation status could be a useful biomarker for cancer immunotherapy response depending on cancer types.

## Introduction

p53 is one of the most important tumor suppressor which is involved in regulation of a wide range of cancer-associated pathways, such as cell cycle, apoptosis, DNA damage repair, metabolism, immune and inflammation, angiogenesis, and metastasis (1). Its encoding gene *TP53* is the most frequently-mutated gene in human cancers and the *TP53* mutation is often associated with a worse prognosis in cancer (2). A number of studies have suggested that p53 plays an important role in tumor immune regulation (3-11). Notably, most of these prior studies consistently showed that p53 played a positive role in antitumor immunosurveillance (3-8, 10, 11). As a result, *TP53* mutations often inhibited antitumor immunity and cancer immunotherapy response (7, 10). However, a few studies have reported that *TP53* mutations could promote antitumor immune activity and immunotherapy response (9, 11, 12). These contradictory findings suggest that the correlation between *TP53* mutations and tumor immunity could be cancer type dependent.

p53 plays a key role in maintaining chromosomal/genomic stability (1). Accordingly, *TP53* mutations may result in chromosomal/genomic instability that contributes to increased tumor mutation burden (TMB) and tumor aneuploidy level (TAL) (1). It has been shown that TMB tends to have a positive correlation with tumor immunity and immunotherapy response (13, 14) and TAL has a negative correlation with them (15). Thus, the alterations of tumor immunity and immunotherapy response could be a consequence of the joint effect of the altered TMB and TAL in *TP53*-mutated cancers relative to *TP53*-wildtype cancers. To prove this hypothesis, we analyzed five cancer cohorts from The Cancer Genome Atlas (TCGA) project (https://portal.gdc.cancer.gov/). The five cancer cohorts included breast invasive carcinoma (BRCA), lung adenocarcinoma (LUAD), stomach adenocarcinoma (STAD), colon adenocarcinoma (COAD), and head and neck squamous cell carcinoma (HNSC). We compared the enrichment levels of diverse immune signatures, TMB, and TAL between *TP53*-mutated and *TP53*-wildtype cancers and analyzed the correlations of TMB and TAL with immune signatures in each of the five cancer cohorts. Furthermore, we compared the immunotherapy response rates between *TP53*-mutated and *TP53*-wildtype cancers in a LUAD cohort (Dong cohort (9)) receiving anti-PD-1/PD-L1 immunotherapy. Moreover, we compared overall survival (OS) prognosis between *TP53*-mutated and *TP53*-wildtype cancers in the immunotherapy setting (Samstein cohort (13)) and in the non-immunotherapy setting (TCGA cohorts). This study revealed the mechanism underlying the distinct correlations of *TP53* mutations with tumor immunity among different caner types.

## Materials and Methods

### Materials

The multi-omics (gene expression and gene somatic mutations) and clinical datasets for six TCGA cancer cohorts (BRCA, LUAD, STAD, COAD, HNSC, and lung squamous-cell carcinoma (LUSC)) were downloaded from the genomic data commons data portal (https://portal.gdc.cancer.gov/). The datasets for two cancer cohorts (Dong cohort (9) and Samstein cohort (13)) with anti-PD-1/PD-L1/CTLA-4 immunotherapy response-associated clinical data were obtained from their associated publications.

### Comparisons of the enrichment levels of immune signatures between *TP53*-mutated and *TP53*-wildtype cancers

We quantified the enrichment level of an immune signature in a tumor sample by the single-sample gene-set enrichment analysis (ssGSEA) score (16) of the immune signature marker gene set. We analyzed three immune signatures, including CD8+ T cells (marker gene *CD8A*), NK cells (marker genes *KLRC1* and *KLRF1*), and immune cytolytic activity (marker genes *GZMA* and *PRF1*). We compared the immune signature enrichment (ISE) levels between *TP53*-mutated and *TP53*-wildtype cancers using Mann-Whitney U test. In addition, we compared the ratios of immune-stimulatory signatures/immune-inhibitory signatures (CD8+/CD4+ regulatory T cells, pro-/anti-inflammatory cytokines, and M1/M2 macrophages) between *TP53*-mutated and *TP53*-wildtype cancers using Mann-Whitney U test. The ratios were the mean expression levels of immune-stimulatory signature marker genes divided by the mean expression levels of immune-inhibitory signature marker genes. Supplementary Table S1 presents the marker genes of these immune signatures.

### Comparisons of TMB and TAL between *TP53*-mutated and *TP53*-wildtype cancers

For each tumor sample, we defined its TMB as the total number of somatic mutations in the tumor and TAL as the ploidy score which was assessed by the ABSOLUTE algorithm (17). We compared TMB and TAL between *TP53*-mutated and *TP53*-wildtype cancers using Mann-Whitney U test.

### Prediction of immune signature enrichment levels using TMB and TAL

We used logistic regression to evaluate the contributions of TMB and TAL in predicting CD8+ T cell enrichment levels in cancer. The predictor TMB was log10 transformed and z-score standardized and TAL was z-score standardized. The CD8+ T cell enrichment level (high (upper third) versus low (bottom third)) was predicted.

### Comparisons of the enrichment levels of immune signatures between high-TMB and low-TMB cancers and between high-TAL and low-TAL cancers

We compared the ISE levels between high-TMB (> median TMB) and low-TMB (< median TMB) cancers and between high-TAL (> median TAL) and low-TAL (< median TAL) cancers using Mann-Whitney U test.

### Survival analyses

We compared OS prognosis between *TP53*-mutated and *TP53*-wildtype cancers in six TCGA cancer cohorts and two immunotherapy-associated cancer cohorts. We used Kaplan-Meier survival curves to show the OS time differences and the log-rank test to assess the significance of OS time differences.

## Results

### Correlation between *TP53* mutations and tumor immunity

Interestingly, we found that the immune signatures (CD8+ T cells, NK cells, and immune cytolytic activity) exhibited significantly higher enrichment levels in *TP53*-mutated cancers than in *TP53*-wildtype cancers in BRCA and LUAD (Mann-Whitney U test, *P*<0.01) (Figure 1A). However, almost all these immune signatures showed significantly lower enrichment levels in *TP53*-mutated cancers than in *TP53*-wildtype cancers in STAD, COAD, and HNSC (Mann-Whitney U test, *P*<0.05) (Figure 1A). Moreover, *TP53*-mutated cancers tended to have higher ratios of immune-stimulatory signatures/immune-inhibitory signatures (CD8+/CD4+ regulatory T cells, pro-/anti-inflammatory cytokines, and M1/M2 macrophages) than *TP53*-wildtype cancers in BRCA and LUAD (Mann-Whitney U test, *P*<0.05) (Figure 1B). In contrast, these ratios were likely lower in *TP53*-mutated cancers than in *TP53*-wildtype cancers in STAD, COAD, and HNSC (Figure 1B). Altogether, these results indicate that the correlation between *TP53* mutations and tumor immunity varies depending on cancer types.

**Figure 1.**
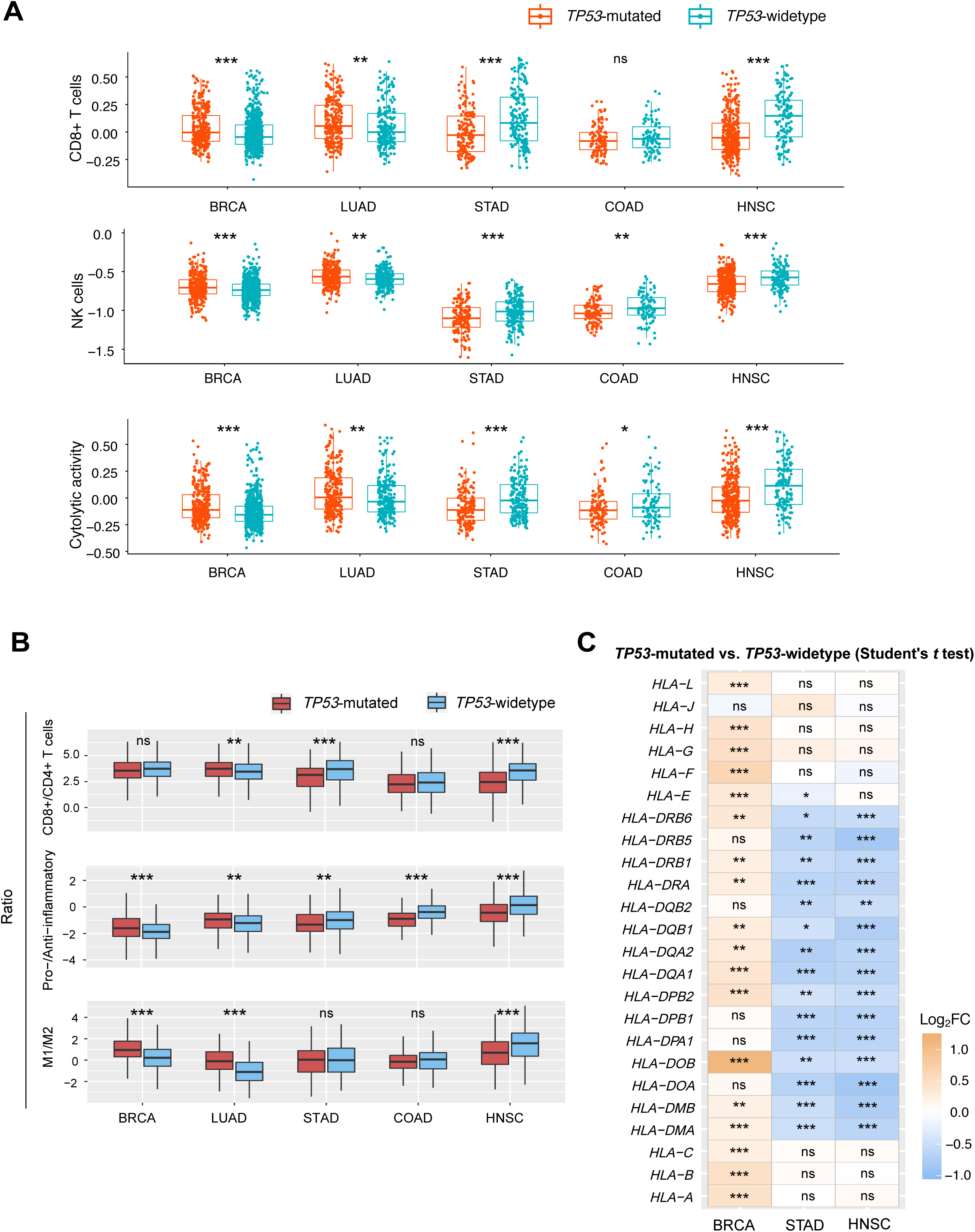
Correlation between *TP53* mutations and tumor immunity. **A.** Three immune signatures (CD8+ T cells, NK cells, and immune cytolytic activity) show significantly higher enrichment levels (ssGSEA scores) in *TP53*-mutated cancers than in *TP53*-wildtype cancers in BRCA and LUAD and lower enrichment levels in *TP53*-mutated cancers than in *TP53*-wildtype cancers in STAD, COAD, and HNSC (Mann-Whitney U test, *P*<0.05). **B.** The ratios of immune-stimulatory signatures/immune-inhibitory signatures (CD8+/CD4+ regulatory T cells, pro-/anti-inflammatory cytokines, and M1/M2 macrophages) tend to be higher in *TP53*-mutated cancers than in *TP53*-wildtype cancers in BRCA and LUAD and to be lower in *TP53*-mutated cancers than in *TP53*-wildtype cancers in STAD, COAD, and HNSC (Mann-Whitney U test, *P*<0.05). **C.** Many HLA genes have higher expression levels in *TP53*-mutated cancers than in *TP53*-wildtype cancers in BRCA and LUAD and have lower expression levels in *TP53*-mutated cancers than in *TP53*-wildtype cancers in STAD, COAD, and HNSC (Student’s *t* test, *P*<0.05). ssGSEA: sample gene-set enrichment analysis (16). BRCA: breast invasive carcinoma. LUAD: lung adenocarcinoma. STAD: stomach adenocarcinoma. COAD: colon adenocarcinoma. HNSC: head and neck squamous cell carcinoma. HLA: human leukocyte antigen. ns: not significant. FC: fold change of mean gene expression levels (*TP53*-mutated cancers/*TP53*-wildtype cancers). * *P<*0.05, ** *P<*0.01, *** *P<*0.001. They also apply to following figures.

Furthermore, we found that numerous human leukocyte antigen (HLA) genes were significantly upregulated in *TP53*-mutated cancers relative to *TP53*-wildtype cancers in BRCA and LUAD (Student’s *t* test, *P*<0.05) (Figure 1C). In contrast, many HLA genes were significantly downregulated in *TP53*-mutated cancers versus *TP53*-wildtype cancers in STAD, COAD, and HNSC (Figure 1C). These results again indicate that the correlation between *TP53* mutations and tumor immunity is cancer type dependent.

### Correlations of *TP53* mutations with TMB and TAL

We compared TMB and TAL between *TP53*-mutated and *TP53*-wildtype cancers in the TCGA cancer cohorts. As expected, both TMB and TAL were significantly higher in *TP53*-mutated cancers than in *TP53*-wildtype cancers except that TMB was lower in *TP53*-mutated COAD compared to *TP53*-wildtype COAD (Mann-Whitney U test, *P*<0.05) (Figure 2). The elevated TMB and TAL could be attributed to the increased chromosomal/genomic instability resulting from p53 inactivation in *TP53*-mutated cancers.

**Figure 2.**
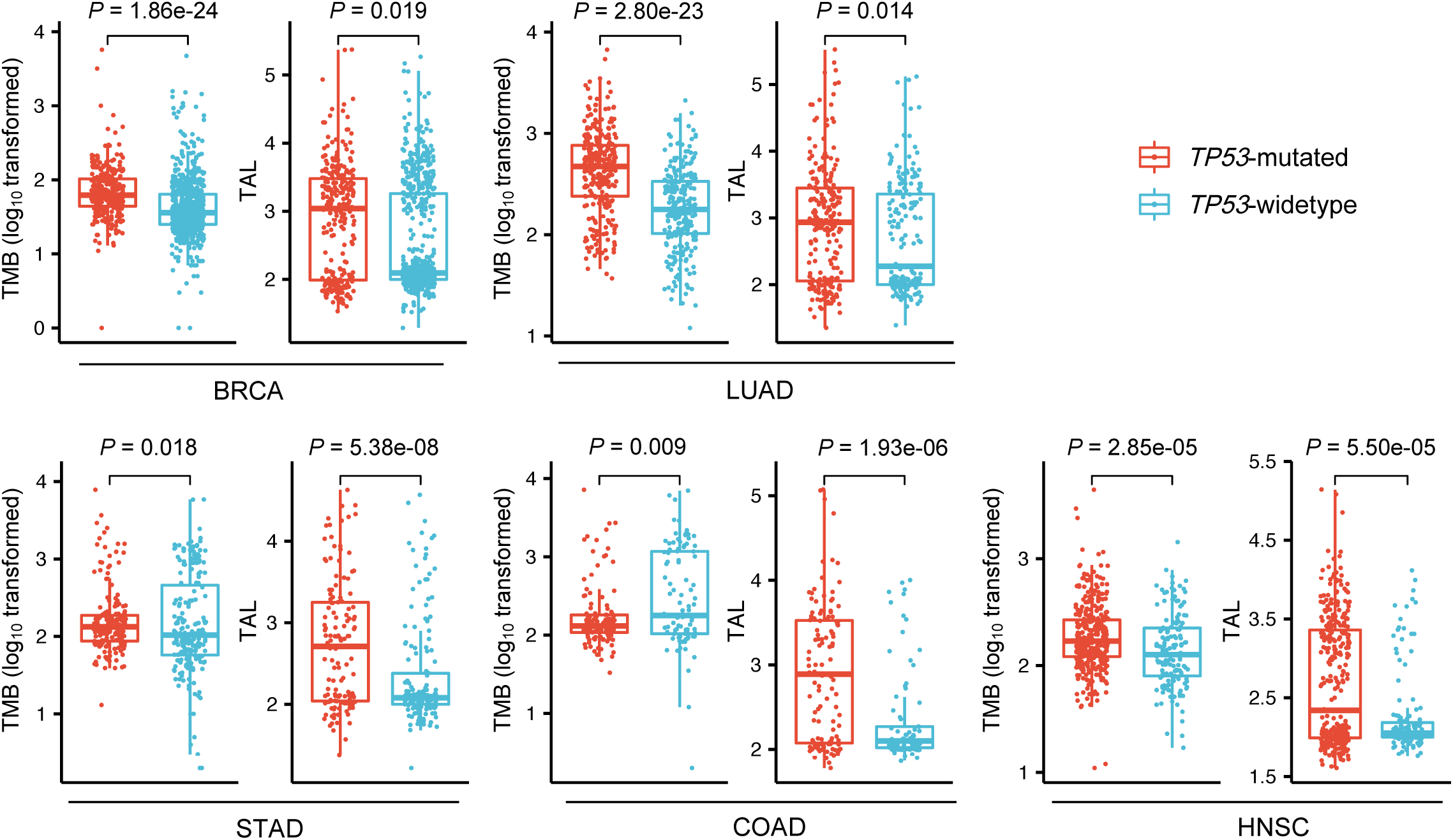
Comparisons of TMB and TAL between *TP53*-mutated and *TP53*-wildtype cancers. Both TMB and TAL are significantly higher in *TP53*-mutated cancers than in *TP53*-wildtype cancers except that TMB is lower in *TP53*-mutated COAD than in *TP53*-wildtype COAD (Mann-Whitney U test, *P*<0.05). The TMB differences are much more marked between *TP53*-mutated and *TP53*-wildtype cancers compared to the TAL differences in BRCA and LUAD while the TAL differences are much more significant in STAD and COAD (a smaller *P* value indicates a more significant difference). TMB: tumor mutation burden. TAL: tumor aneuploidy level. They also apply to following figures.

Interestingly, we found that the TMB differences were much more marked between *TP53*-mutated and *TP53*-wildtype cancers compared to the TAL differences in BRCA and LUAD while the TAL differences were much more significant in STAD and COAD (Figure 2). It indicates that the altered degree of TMB and TAL resulting from *TP53* mutations depends on cellular context. This finding supports the notion that p53 action and regulation are context-dependent (1).

### Correlations of TMB and TAL with tumor immunity

We compared the enrichment levels of immune signatures between high-TMB and low-TMB cancers in each of the five TCGA cancer cohorts and found that these immune signatures were more highly enriched in high-TMB cancers than in low-TMB cancers (Mann-Whitney U test, *P*<0.05) (Figure 3A). The ratios of immune-stimulatory signatures/immune-inhibitory signatures were higher in high-TMB cancers than in low-TMB cancers (Mann-Whitney U test, *P*<0.05) (Figure 3A). Moreover, we compared the enrichment levels of immune signatures between high-TAL and low-TAL cancers in each of the five TCGA cancer cohorts. We found that the immune signatures were more lowly enriched in high-TAL cancers than in low-TAL cancers (Mann-Whitney U test, *P*<0.05) (Figure 3A). The ratios of immunostimulatory signature scores versus immunoinhibitory signature scores were lower in high-TAL cancers than in low-TAL cancers (Mann-Whitney U test, *P*<0.05) (Figure 3A). A number of HLA genes exhibited significantly higher expression levels in high-TMB cancers than in low-TMB cancers while significantly lower expression levels in high-TAL cancers than in low-TAL cancers (Student’s *t* test, *P*<0.05) (Supplementary Figure S1). These results were consistent with previous observation that tumor immunity had a positive correlation with TMB and an inverse correlation with TAL (15).

**Figure 3.**
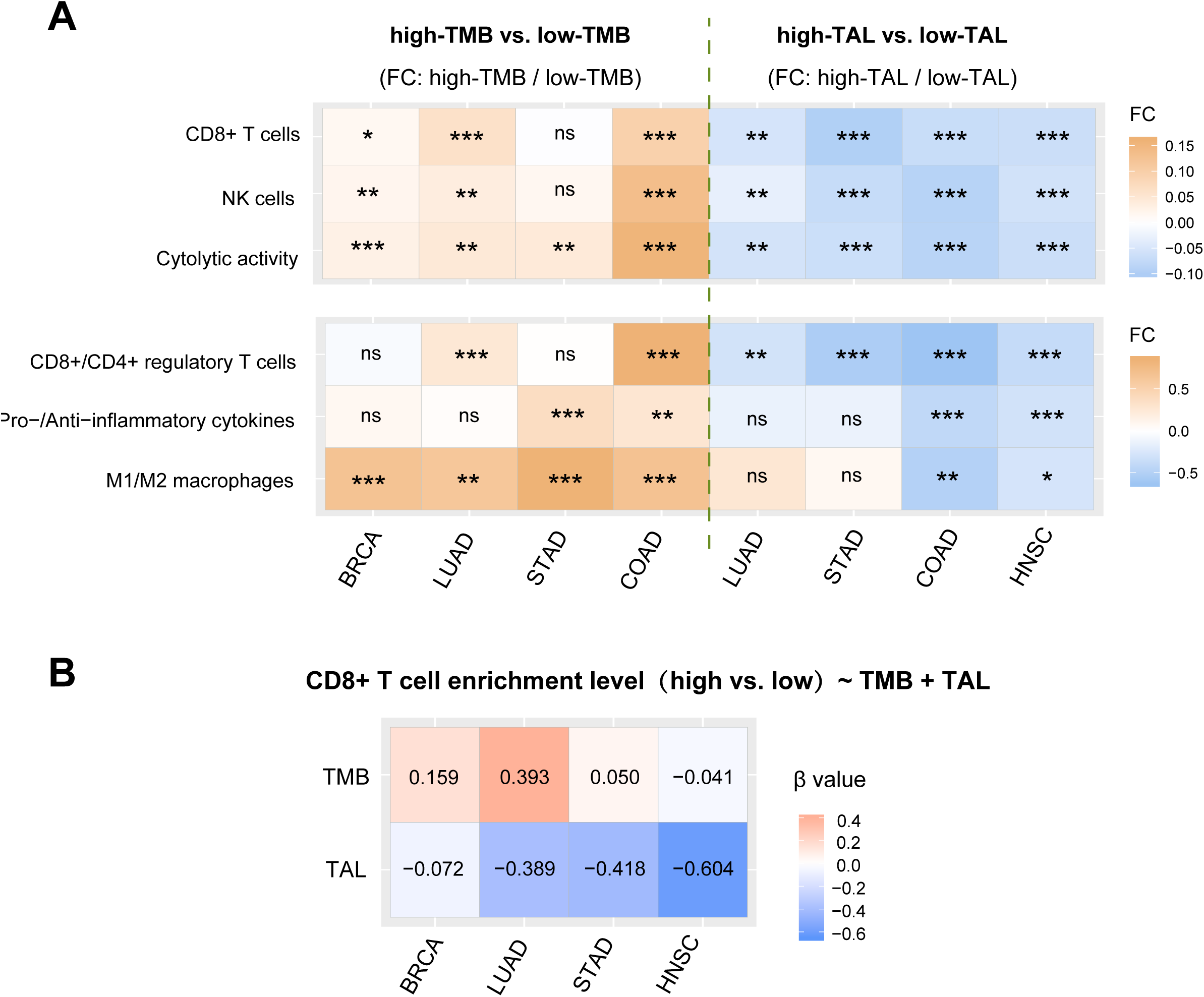
Correlations of TMB and TAL with tumor immunity. **A.** The immune signature enrichment levels and the ratios of immune-stimulatory signatures/immune-inhibitory signatures are higher in high-TMB (TMB > median) cancers than in low-TMB (TMB < median) cancers and are lower in high-TAL (TAL > median) cancers than in low-TAL (TAL < median) cancers (Mann-Whitney U test, *P*<0.05). FC: fold change of mean immune signature enrichment levels or ratios. **B.** Logistic regression analysis shows that TMB is a stronger predictor than TAL in predicting CD8+ T cell enrichment levels in BRCA and LUAD and TAL a stronger predictor than TMB in predicting CD8+ T cell enrichment levels in STAD and HNSC.

Furthermore, we used the logistic regression model with two predictors (TMB and TAL) to predict CD8+ T cell enrichment levels in each of the five TCGA cancer cohorts. We found that TMB was a stronger predictor than TAL in predicting CD8+ T cell enrichment levels in BRCA (TMB: β=0.159, *P*=0.050 versus TAL: β=-0.072, *P*=0.422) as well as in LUAD (TMB: β=0.393, *P*=0.002 versus TAL: β=-0.389, *P*=0.002) (Figure 3B). In contrast, TAL was a stronger predictor than TMB in predicting CD8+ T cell enrichment levels in STAD (TMB: β=0.050, *P*=0.742 versus TAL: β=-0.418, *P*=0.007) and in HNSC (TMB: β=-0.041, *P*=0.732 versus TAL: β=-0.604, *P*=2.25×10^-6^) (Figure 3C). These results indicate that TMB affects ISE to a greater degree than TAL does in BRCA and LUAD while TAL affects ISE more highly than TMB in STAD and HNSC. Since TMB and TAL are positively and negatively associated with tumor immunity, respectively, and *TP53* mutations enhance both TMB and TAL, *TP53* mutations may result in increased or reduced tumor immunity depending on which of TMB and TAL has a stronger impact on tumor immunity. It could explain why *TP53*-mutated cancers have higher tumor immunity than *TP53*-wildtype cancers in BRCA and LUAD while *TP53*-mutated cancers have lower tumor immunity than *TP53*-wildtype cancers in STAD and HNSC.

### Correlations of *TP53* mutations with survival prognosis in cancer

*TP53* mutations have been associated with a worse survival in various cancers (2). However, *TP53* mutations showed no a significant correlation with OS in BRCA (log-rank test, *P*=0.162). A possible explanation is that *TP53* mutations promote antitumor immune response in BRCA (Figure 1A). In addition, we found that *TP53* mutations were significantly associated with a worse OS in LUAD and HNSC (log-rank test, *P*<0.05) but showed no a significant correlation with OS in STAD, COAD, and gastrointestinal (GI) cancers. Furthermore, we examined the association between *TP53* mutations and OS in these cancer types in a pan-cancer cohort (Samstein cohort (13)) receiving anti-PD-1/PD-L1/CTLA-4 immunotherapy. We found that *TP53* mutations were significantly associated with a worse OS in COAD and GI cancers (log-rank test, *P*<0.05) and exhibited no a significant correlation with OS in BRCA, LUAD, STAD, and HNSC in Samstein cohort. The negative correlation between *TP53* mutations and OS in COAD and GI cancers in the immunotherapy setting rather than in the non-immunotherapy setting (Figure 4A) could be attributed to the better immunotherapy response in the *TP53*-wildtype subtype than in the *TP53*-mutated subtype of COAD and GI cancers. On the other hand, the negative correlation between *TP53* mutations and OS in LUAD in the non-immunotherapy setting rather than in the immunotherapy setting (Figure 4A) could be a consequence of the better immunotherapy response in *TP53*-mutated LUAD than in *TP53*-wildtype LUAD. This speculation was confirmed in a LUAD cohort (Dong cohort (9)) receiving anti-PD-1/PD-L1 immunotherapy where *TP53*-mutated LUAD had a 1.5 times higher immunotherapy response rate (57.1%) than *TP53*-wildtype LUAD (37.5%) and *TP53*-mutated LUAD showed a better OS than *TP53*-widtype LUAD (9). These results suggest that the association between *TP53* mutations and immunotherapy response is cancer type dependent. This conclusion can be evidenced by two recent studies showing that *TP53* mutations could promote immunotherapy response in LUAD (9) and inhibit immunotherapy response in melanoma (10). Furthermore, we found that *PD-L1* expression levels exhibited a significant positive correlation with *TP53* mutations in BRCA and LUAD while a significant inverse correlation with *TP53* mutations in STAD, COAD, and HNSC (Student’s *t* test, *P*<0.05) (Figure 4B). This could explain why *TP53*-mutated LUAD had a better immunotherapy response than *TP53*-wildtype LUAD and *TP53*-mutated COAD had a worse immunotherapy response than *TP53*-wildtype COAD since PD-L1 expression is a biomarker for anti-PD-1/PD-L1 immunotherapy response in cancer (18).

**Figure 4.**
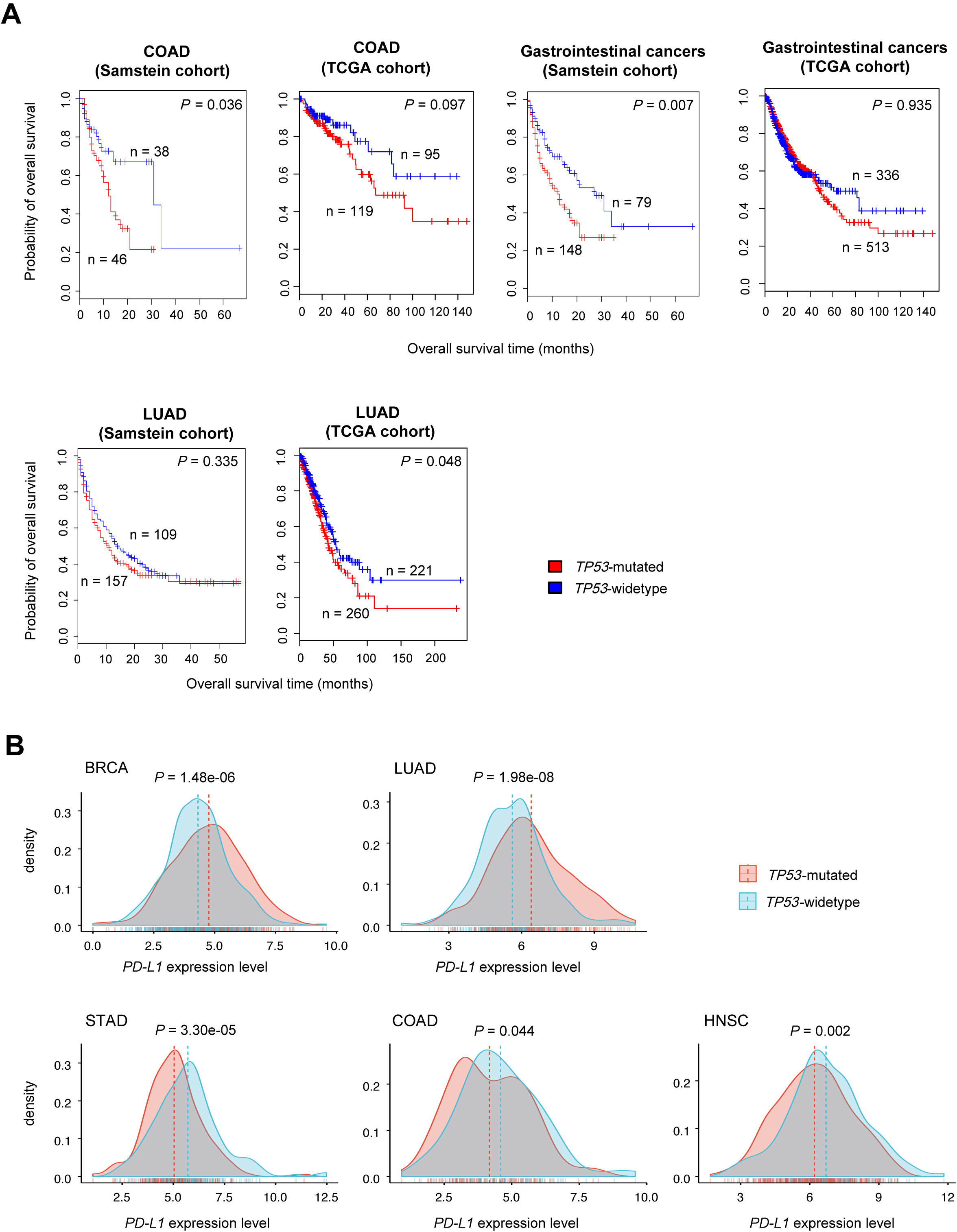
Correlations of *TP53* mutations with survival prognosis in cancer. A. Kaplan-Meier survival curves show that *TP53* mutations have a negative correlation with overall survival (OS) in COAD and gastrointestinal cancers in the immunotherapy setting (Samstein cohort (13)) but have no a significant correlation with OS in the non-immunotherapy setting (TCGA cohorts) and that *TP53* mutations have a negative correlation with OS in LUAD in the non-immunotherapy setting but have no a significant correlation with OS in the immunotherapy setting. Log-rank test *P* values are shown. **B.** *PD-L1* expression levels are significantly higher in *TP53*-mutated cancers than in *TP53*-wildtype cancers in BRCA and LUAD and are significantly lower in *TP53*-mutated cancers than in *TP53*-wildtype cancers in STAD, COAD, and HNSC (Student’s *t* test, *P*<0.05).

## Discussion

A number of studies have revealed that cellular context is essential to p53 function and activity (1, 19). In this study, we demonstrated that regulation of tumor immunity by p53 was cancer type dependent. We found that the correlation between *TP53* mutations and tumor immunity was positive in BRCA and LUAD while was negative in STAD, COAD, and HNSC. To our knowledge, this is the first study showing distinct correlations of *TP53* mutations with tumor immunity in different cancer types. Interestingly, we found that even in the cancers originating from the same organ, the correlations of *TP53* mutations with tumor immunity could be distinct. For example, the correlation between *TP53* mutations and tumor immunity tended to be negative in LUSC (Supplementary Figure S2A) versus the positive correlation between *TP53* mutations and tumor immunity in LUAD (Figure 1A). Moreover, *TP53* mutations were associated with a worse OS in LUSC in the immunotherapy setting (Samstein cohort) (log-rank test, *P*=0.046) (Supplementary Figure S2B) while showed no a significant association with OS in LUAD in this cohort (log-rank test, *P*=0.335) (Figure 4A).

p53 is a significant protector of genome stability and *TP53* mutations often comprise the p53 function in maintaining genome stability (20). As a result, *TP53*-mutated cancers tend to exhibit higher TMB and TAL than *TP53*-wildtype cancers (Figure 2) since genome instability leads to increased TMB and TAL in cancer. An exception was COAD in which *TP53*-mutated cancers had lower TMB and higher TAL than *TP53*-wildtype cancers (Figure 2). If the genome instability caused by *TP53* mutations is restricted to local and small DNA sequences, such as microsatellite instability/deficient mismatch repair (MSI/dMMR), a higher degree of TMB differences would be presented between *TP53*-mutated and *TP53*-wildtype cancers. In contrast, if the genome instability caused by *TP53* mutations is the alterations of chromosomal or large DNA sequences, such as DNA ploidy or copy number alterations, a higher degree of TAL differences would be shown between *TP53*-mutated and *TP53*-wildtype cancers. Because TMB and TAL have a positive and a negative correlation with tumor immunity, respectively, *TP53* mutations may increase or reduce tumor immunity depending on the counteractive effect of TMB and TAL on tumor immunity (Figure 5). We confirmed this hypothesis by analyzing five TCGA cancer cohorts. We further demonstrated that the *TP53* mutation status could be a positive or a negative predictor for cancer immunotherapy response depending on cancer types.

**Figure 5.**
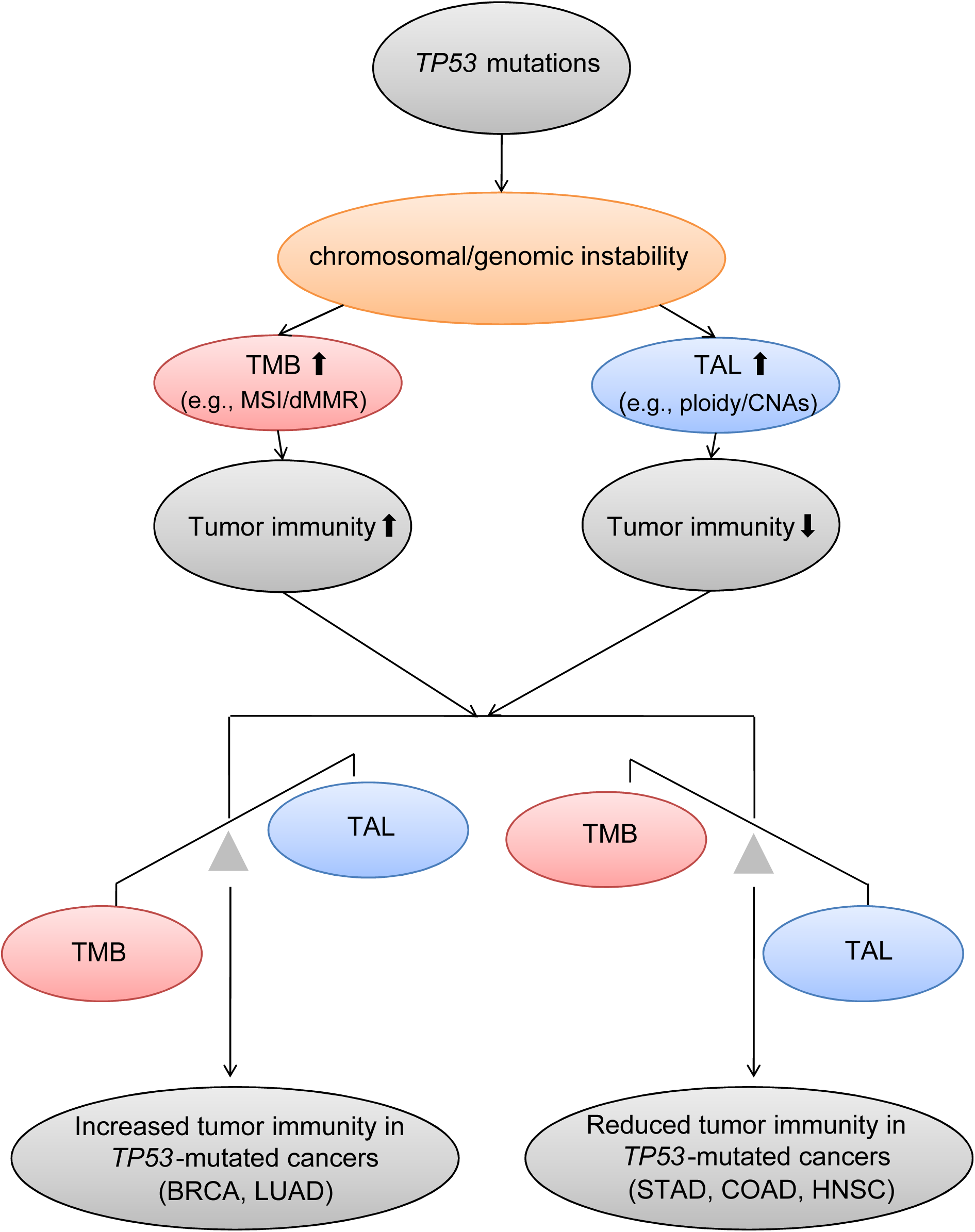
The mechanism underlying the distinct correlations of *TP53* mutations with tumor immunity among different caner types. MSI: microsatellite instability. dMMR: deficient mismatch repair. CNAs: copy number alterations.

## Conclusions

The correlations between *TP53* mutations and tumor immunity and immunotherapy response are cancer type dependent. These findings have potential clinical implications for cancer immunotherapy.

## Disclosure of Potential Conflicts of Interest

The authors declare that they have no competing interests.

## Authors’ Contributions

### Conception and design

X. Wang

### Development of methodology

X. Wang, L. Li, M. Li

### cquisition of data (provided animals, acquired and managed patients, provided facilities, etc.)

L. Li, M. Li

### Analysis and interpretation of data (e.g., statistical analysis, biostatistics, computational analysis)

L. Li, M. Li, X. Wang

### Writing, review, and/or revision of the manuscript

X. Wang

### Administrative, technical, or material support (i.e., reporting or organizing data, constructing databases)

L. Li, M. Li

### Study supervision

X. Wang

## Acknowledgments

We thank Mr. Zehang Jiang from China Pharmaceutical University for help in data analyses.

## Supplementary data

**Supplementary Table S1. The gene sets that represent different immune signatures.**

**Supplementary Figure S1.** A number of HLA genes show significantly higher expression levels in high-TMB cancers than in low-TMB cancers and significantly lower expression levels in high-TAL cancers than in low-TAL cancers (Student’s *t* test, *P*<0.05).

**Supplementary Figure S2. Correlations of *TP53* mutations with tumor immunity and overall survival (OS) prognosis in LUSC. A.** The correlation between *TP53* mutations and tumor immunity tends to be negative in LUSC. **B.** Kaplan-Meier survival curves show that *TP53* mutations are associated with a worse OS in LUSC in the immunotherapy setting (Samstein cohort (13)) (log-rank test, *P*=0.046) and are associated with a better OS in LUSC in the non-immunotherapy setting (TCGA cohort). LUSC: lung squamous-cell carcinoma.

## References

1. Kastenhuber ER & Lowe SW (2017) Putting p53 in Context. Cell 170(6):1062–1078.

2. Wang X & Sun Q (2017) TP53 mutations, expression and interaction networks in human cancers. Oncotarget 8(1):624–643.

3. Zitvogel L & Kroemer G (2015) CANCER. A p53-regulated immune checkpoint relevant to cancer. Science 349(6247):476–477.

4. Textor S, et al. (2011) Human NK cells are alerted to induction of p53 in cancer cells by upregulation of the NKG2D ligands ULBP1 and ULBP2. Cancer research 71(18):5998–6009.

5. Shatz M, Menendez D, & Resnick MA (2012) The human TLR innate immune gene family is differentially influenced by DNA stress and p53 status in cancer cells. Cancer research 72(16):3948–3957.

6. Guo G, Yu M, Xiao W, Celis E, & Cui Y (2017) Local activation of p53 in the tumor microenvironment overcomes immune suppression and enhances antitumor immunity. Cancer research.

7. Jiang Z, Liu Z, Li M, Chen C, & Wang X (2018) Immunogenomics Analysis Reveals that TP53 Mutations Inhibit Tumor Immunity in Gastric Cancer. Translational oncology 11(5):1171–1187.

8. Wang B, Niu D, Lai L, & Ren EC (2013) p53 increases MHC class I expression by upregulating the endoplasmic reticulum aminopeptidase ERAP1. Nature communications 4:2359.

9. Dong ZY, et al. (2017) Potential Predictive Value of TP53 and KRAS Mutation Status for Response to PD-1 Blockade Immunotherapy in Lung Adenocarcinoma. Clinical cancer research: an official journal of the American Association for Cancer Research 23(12):3012–3024.

10. Xiao W, et al. (2018) TP53 Mutation as Potential Negative Predictor for Response of Anti-CTLA-4 Therapy in Metastatic Melanoma. EBioMedicine 32:119–124.

11. Ham SW, et al. (2019) TP53 gain-of-function mutation promotes inflammation in glioblastoma. Cell Death Differ 26(3):409–425.

12. Liu Z, et al. (2019) TP53 Mutations Promote Immunogenic Activity in Breast Cancer. Journal of Oncology 2019(Article ID 5952836):1–19.

13. Samstein RM, et al. (2019) Tumor mutational load predicts survival after immunotherapy across multiple cancer types. Nature genetics 51(2):202–206.

14. Goodman AM, et al. (2017) Tumor Mutational Burden as an Independent Predictor of Response to Immunotherapy in Diverse Cancers. Molecular cancer therapeutics 16(11):2598–2608.

15. Davoli T, Uno H, Wooten EC, & Elledge SJ (2017) Tumor aneuploidy correlates with markers of immune evasion and with reduced response to immunotherapy. Science 355(6322).

16. Hanzelmann S, Castelo R, & Guinney J (2013) GSVA: gene set variation analysis for microarray and RNA-seq data. BMC bioinformatics 14:7.

17. Carter SL, et al. (2012) Absolute quantification of somatic DNA alterations in human cancer. Nature biotechnology 30(5):413–421.

18. Patel SP & Kurzrock R (2015) PD-L1 Expression as a Predictive Biomarker in Cancer Immunotherapy. Molecular cancer therapeutics 14(4):847–856.

19. Andrysik Z, et al. (2017) Identification of a core TP53 transcriptional program with highly distributed tumor suppressive activity. Genome research 27(10):1645–1657.

20. Eischen CM (2016) Genome Stability Requires p53. Cold Spring Harbor perspectives in medicine 6(6).

