## Supplementary figures and images for "Cancer Type-Dependent Correlations between *TP53* Mutations and Antitumor Immunity"

### Supplemental Figure 1

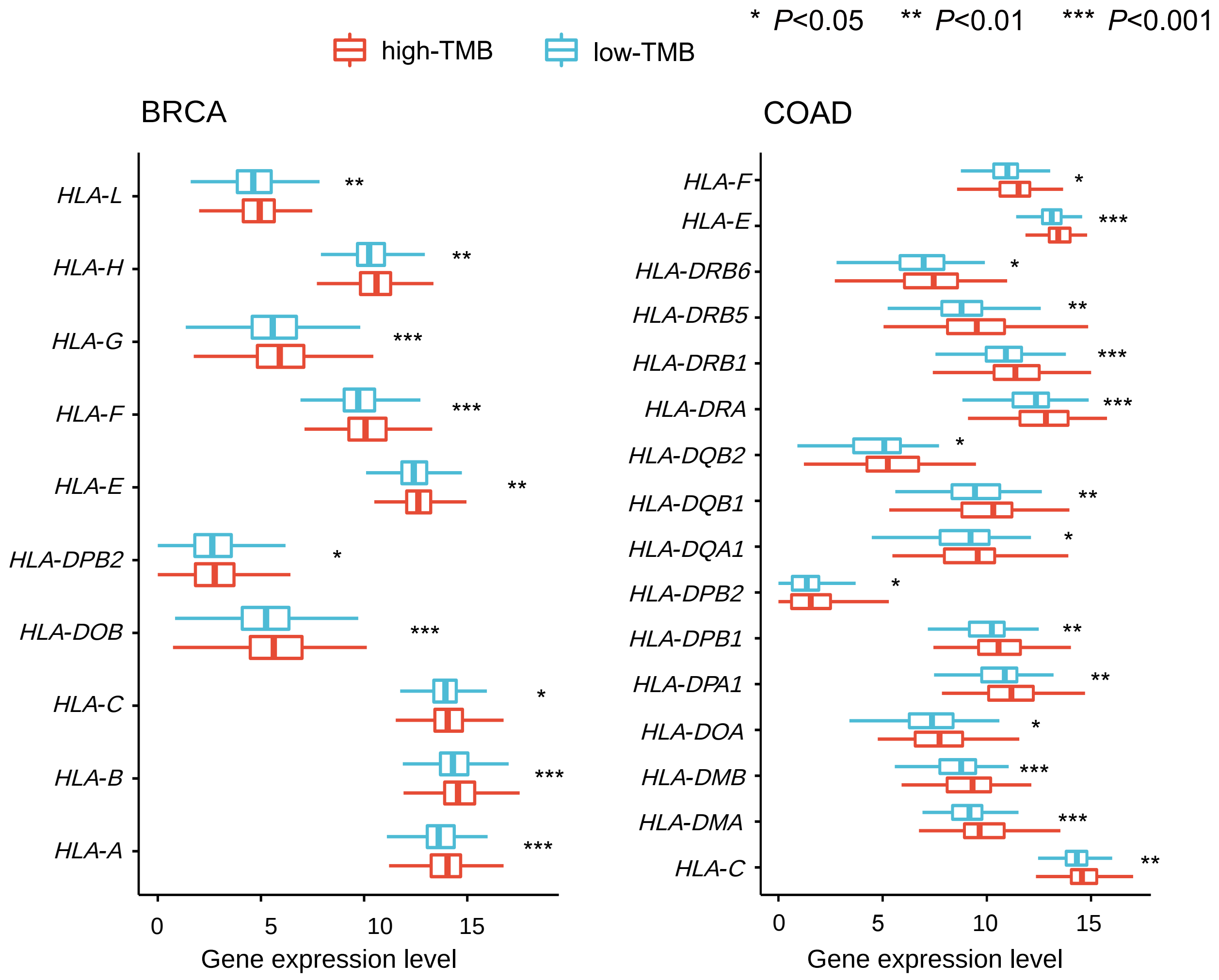

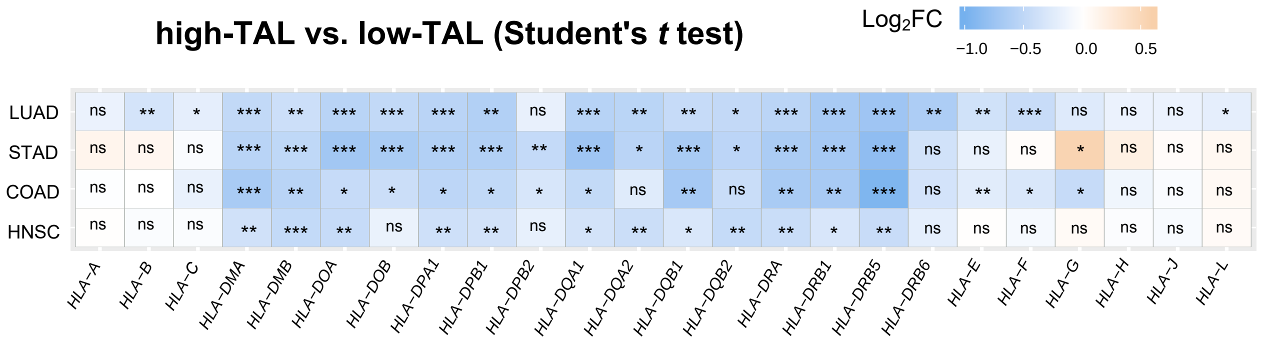

### Supplemental Figure 2

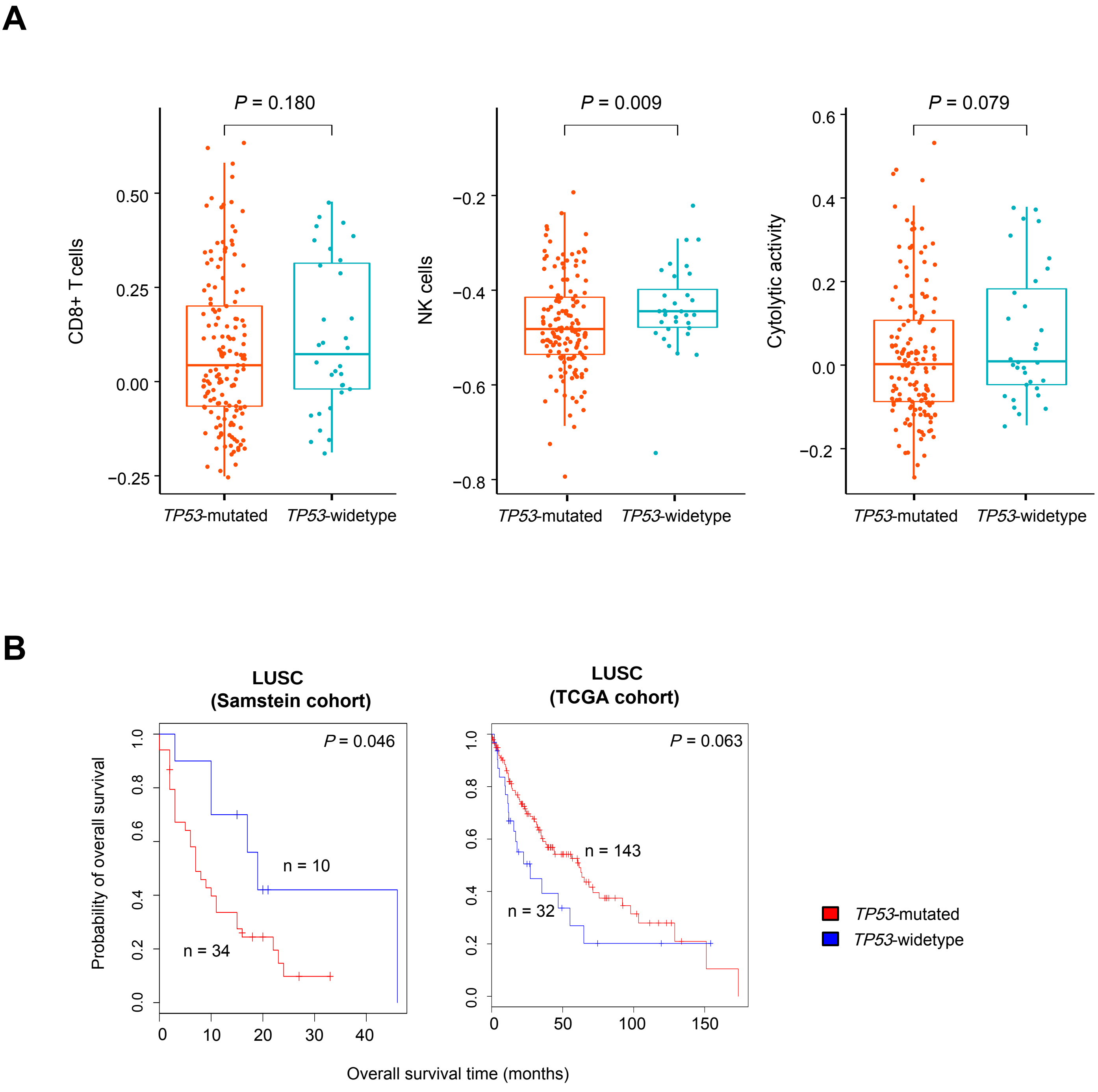
